# Beyond immunity: a transcriptomic landscape of Plasmodium’s modulation of mosquito metabolic pathways

**DOI:** 10.1101/2024.09.09.611609

**Authors:** L García-Longoria, A Berthomieu, O Hellgren, A Rivero

**Affiliations:** Department of Anatomy, Cell Biology and Zoology. Avenida de Elvas, edificio Margarita Salas CP 06006 Universidad de Extremadura, Badajoz, Spain; MIVEGEC (Univ Montpellier, CNRS, IRD), Montpellier, France; Evolutionary Ecology and Infection biology, Lund University, Department of Biology, Sweden

## Abstract

The focus of mosquito-*Plasmodium* interactions has predominantly been centered on mosquito immunity, revealing key mechanisms by which mosquitoes attempt to combat *Plasmodium* infection. However, recent evidence suggests that beyond immunity, a multitude of mosquito physiological and metabolic pathways play crucial roles in determining whether the parasite completes its development within the mosquito. We review which of these metabolic pathways are potentially modulated by *Plasmodium*, revealing a fragmented and occasionally contradictory state of knowledge. We then present a comprehensive transcriptomic analysis of *Plasmodium*-infected and uninfected mosquitoes, examining gene expression of crucial genes across different stages of the parasite’s development. These genes range from key enzymes and proteins involved in gut structure and function, to genes involved in egg production and resorption, salivary gland invasion and mosquito behaviour. For this purpose, we use a non-model system consisting of the avian malaria parasite *Plasmodium relictum*, an invasive parasite threatening bird biodiversity across the world, and its natural vector, the mosquito *Culex pipiens*. Our results reveal how at each stage of its development within the mosquito, *Plasmodium* modulates a myriad of mosquito metabolic pathways, in ways that potentially favour its survival and the completion of its life cycle. We discuss whether this constitutes sufficient evidence of parasite-driven manipulation or whether the changes are simply the mosquito’s response to the infection, which the parasite may serendipitously exploit to enhance its fitness. Our study extends the comparative transcriptomic analyses of malaria-infected mosquitoes beyond human and rodent parasites, and provides insights into the degree of conservation of metabolic pathways and into the selective pressures exerted by *Plasmodium* parasites on their vectors.

## Introduction

What makes a good malaria vector? This question has been central to malaria control efforts even since Ronald Ross discovered that *Plasmodium* parasites are not transmitted through the foul smells emanating from swamps but through the bites of infected mosquitoes [1]. The ability of mosquitoes to temporarily host, and ultimately transmit, malaria parasites is the outcome of complex co-evolutionary processes in which mosquitoes and parasites with overlapping geographic distributions engage in adaptations and counter-adaptations aimed at maximizing their respective finesses. Disentangling the mechanisms underlying mosquito-*Plasmodium* interactions is crucial not only for comprehending the epidemiology and evolution of the disease but also for developing effective control strategies.

Far from being mere flying syringes, mosquitoes provide a very specific environment in which the malaria parasites differentiate, proliferate, and migrate to the correct tissues to ensure transmission to the next host. The parasite has a complex life cycle within the mosquito, switching between haploid, diploid and syncytial forms, invading drastically different environments (midgut, haemolymph, salivary glands), and undergoing dramatic booms and boosts in population fluctuations [2].

Most research on mosquito-*Plasmodium* interactions has concentrated on mosquito immunity due to its pivotal role in determining the success of malaria transmission [3,4]. As a result, there have been significant advances in our understanding of the cellular and humoral effectors and upstream immune pathways that lead to the lysis, melanization, or phagocytosis of an invading pathogen [3]. The results of studies comparing the immune transcriptome of *Plasmodium*-infected and uninfected mosquitoes strongly suggest that the parasite may be able to manipulate the mosquito immune response to its own advantage [5]. There is increasing evidence, however, that immunity is not the only determinant of the parasite’s fate within the mosquito. A plethora of finely tuned and timed physiological events, ranging from eye pigment precursors, to metabolic enzymes and digestive proteases may be equally, if not more, important in determining the fate of the parasite within the mosquito. Growing evidence indicates that many of these pathways are disrupted during a *Plasmodium* infection, suggesting that the parasite may have evolved strategies to modulate these pathways thereby enhancing and extending the duration of infection and transmission. To provide the necessary context for understanding how the parasite interacts with, and potentially manipulates, various physiological pathways to enhance its survival and transmission, it is essential to briefly explain the life cycle of *Plasmodium* within the mosquito.

The parasite starts its journey inside the mosquito when a female takes a blood meal containing male and female *gametocytes*. Once inside the mosquito midgut, the female gametocyte transforms directly into a single *female gamete*, while the male gametocyte undergoes three rapid cycles of mitosis to generate 8 flagellated *male gametes* [6]. The male and female gametes fuse to form a *zygote*, the only diploid stage of the *Plasmodium* cycle. The processes of meiosis and recombination are still poorly understood, but they take place directly after the formation of the zygote and result in a mobile tetraploid stage called an *ookinete* [7]. This ookinete has to traverse both the peritrophic matrix (PM), a chitinous barrier that protects the gut against the oxidative effects of the digested blood [8,9], and the midgut epithelium before it can attach itself to the basal lamina and develop into an *oocyst*. The oocyst is the longest developmental stage of the *Plasmodium* life cycle, lasting between 10–15 days dependent upon the species. During this time the oocyst will grow in size and simultaneously undergo several rounds of nuclear division resulting in the production of thousands of haploid nuclei. During a massive cytokinesis, thousands of haploid *sporozoites* are released into the haemolymph, from where they migrate to the salivary glands of the mosquito, ready to be transmitted to a new host in the next blood feeding event.

At each stage of its development within the mosquito, *Plasmodium* exploits a myriad of mosquito metabolic pathways, which are crucial for its survival and completion of its life cycle. For instance the gametocyte to gamete transformation is triggered by environmental stimuli, particularly a drop in temperature and an increase in pH [6], but also by the presence of xanthurenic acid (XA), a molecule produced through the tryptophan metabolism (kynurenine) pathway of the mosquito, where it acts as a precursor for its eye pigments [10]. Digestive enzymes such as trypsins, chymotrypsins, and carboxypeptidases, secreted by the mosquito to digest the blood meal, also seem play a crucial role in the parasite’s midgut survival and successful formation of oocysts. Trypsin, for instance, is required to activate the inactive chitinase zymogen secreted by certain *Plasmodium* species to break down and traverse the chitinous PM [11,12]. The massive production of sporozoites is very metabolically demanding, and there is increasing evidence that the parasite is heavily dependent on a panel mosquito lipid [13,14] and sugar [15,16] transport proteins as well as enzymatic [17] and hormonal pathways [18] to provide the developing oocysts with these crucial resources.

Trypsin, XA, carboxypeptidase A, lipid and sugar transport proteins are prime examples of how malaria parasites may have harnessed several of the mosquito’s metabolic pathways for their own benefit, but recent research shows there may be many others. Here, we first carry out a comprehensive review of studies published to date on how mosquito metabolism is modified in a way that potentially enhances the parasite’s fitness during a *Plasmodium* infection. This review reveals a very fragmented, and sometimes contradictory, picture of our current state of knowledge (Tables 1-5). Most of these studies have examined each physiological factor in isolation, using different *Plasmodium*-mosquito combinations, including combinations not found in nature. Consequently, we lack an integrative understanding of how non-immune mosquito physiological pathways may facilitate *Plasmodium* development within the mosquito.

**Table 1:**
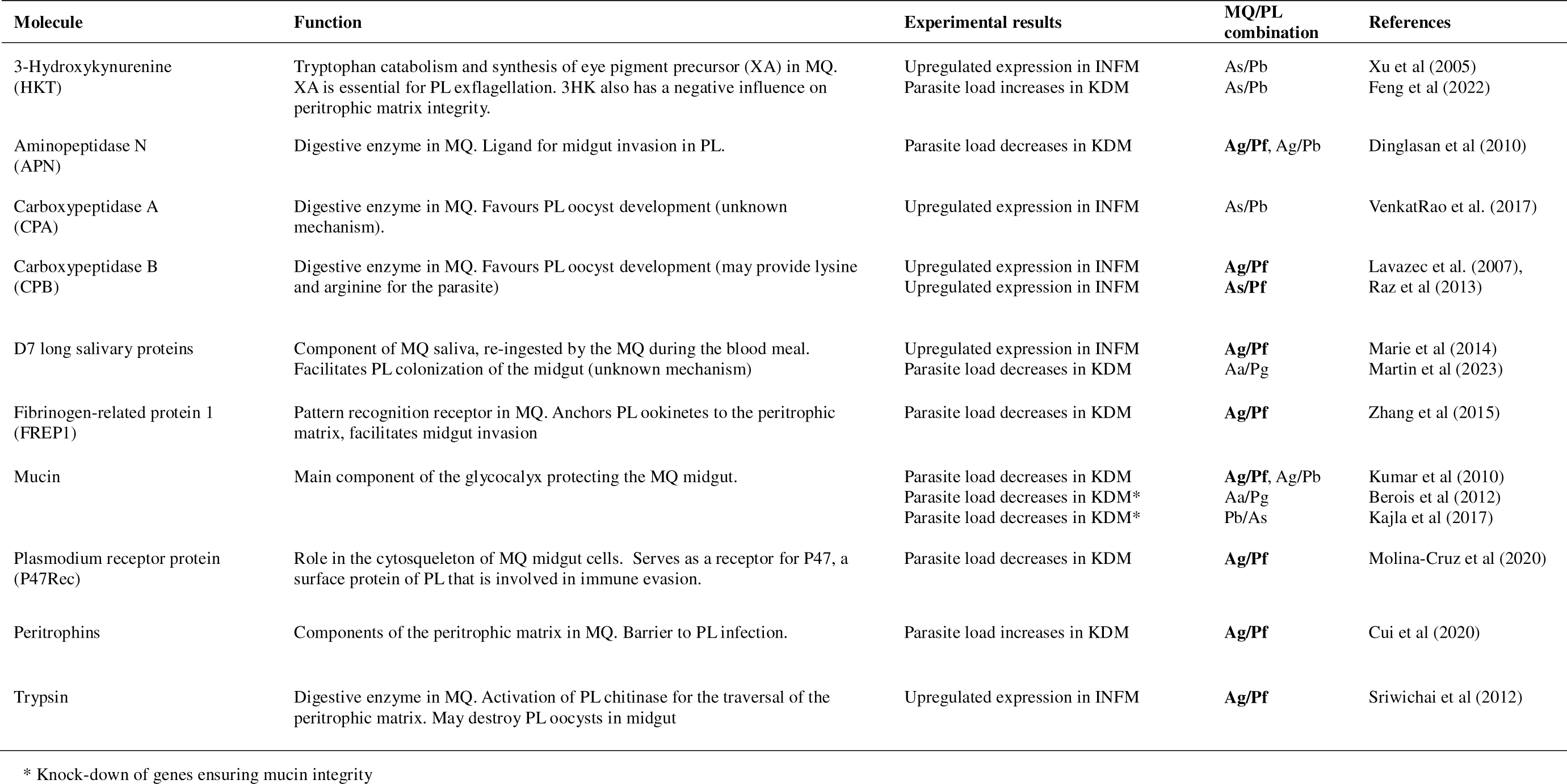
Role of non-immune physiological factors previously reported as being putatively involved on *Plasmodium* invasion and survival in the mosquito midgut. Natural mosquito/Plasmodium combinations are in bold. MQ = mosquito, PL = *Plasmodium*, Ag = *Anopheles gambiae*, Ac = *Anopheles coluzzi*, As = *Anopheles stephensi*, Aa= Aedes aegypti, Pf = *Plasmodium falciparum* (human), Pb = *Plasmodium berguei* (rodent), Pg = *Plasmodium gallinaceum* (avian), INFM = experimentally infected mosquitoes, KDM = knock-down mosquitoes, XA = Xanthurenic Acid.

**Table 2:**
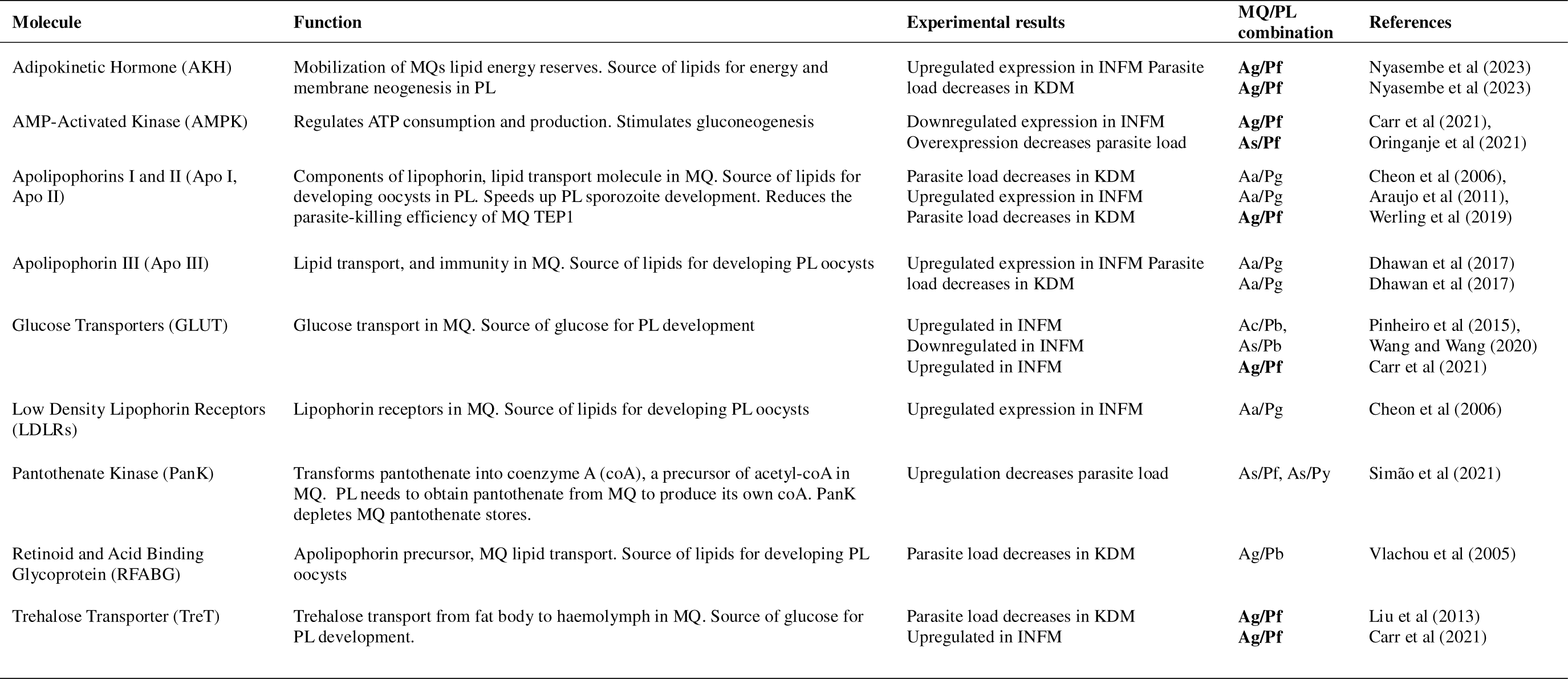
Role of non-immune physiological proteins previously reported as being putatively involved on *Plasmodium* energetic metabolism in the mosquito. Natural mosquito/Plasmodium combinations are in bold. MQ = mosquito, PL = *Plasmodium*, Ag = *Anopheles gambiae*, Ac = *Anopheles coluzzi*, As = *Anopheles stephensi*, Aa= Aedes aegypti, Pf = *Plasmodium falciparum* (human), Pb = *Plasmodium berguei* (rodent), Py = *Plasmodium yoelii* (rodent), Pg = *Plasmodium gallinaceum* (avian), INFM = experimentally infected mosquitoes, KDM = knock-down mosquitoes.

**Table 3:** Role of non-immune physiological proteins previously reported as being putatively involved on mosquito fecundity. Natural mosquito/Plasmodium combinations are in bold. MQ = mosquito, PL = *Plasmodium*, Ag = *Anopheles gambiae*, Ac = *Anopheles coluzzi*, As = *Anopheles stephensi*, Aa= Aedes aegypti, Pf = *Plasmodium falciparum* (human), Pb = *Plasmodium berguei* (rodent), Py = *Plasmodium yoelii* (rodent), Pg = *Plasmodium gallinaceum* (avian), INFM = experimentally infected mosquitoes, KDM = knock-down mosquitoes.

| Molecule | Function | Experimental results | MQ/PL combination | References |
| --- | --- | --- | --- | --- |
| 20-Hydroxyecdysone (20E) | Reproductive hormone, involved in MQ egg production. Stimulates Vg synthesis. | Downregulated expression in INFM<br>Downregulation decreases parasite load | Aa/Pg<br>**Ag/Pf** | Araujo et al (2011)<br>Werling et al (2019) |
| Allatoicase (ALLC) | Converts excess uric acid (antioxidant) into urea in MQ. Contributes to redox homeostasis in MQ. A reduction in uric acid may be correlated to a reduction of fecundity in MQ. | Upregulated expression in INFM | Ag/Pb | Kwon and Smith (2022) |
| Branched chain amino acid transferase (BCAT) | Involved in egg production in MQ | Upregulation decreases parasite load | **Ac/Pf** | Lampe et al (2019) |
| Nutrient sensing pathway (TOR) | Source of lipids for energy and membrane neogenesis in MQ. Regulates cellular metabolism. Conversion of blood meal into eggs. | Downregulated expression in INFM | **Ag/Pf** | Carr et al (2021) |
| Urate Oxidase (UO) | Converts excess uric acid (antioxidant) into urea in MQ. Contributes to redox homeostasis in MQ. A reduction in uric acid may be correlated to a reduction of fecundity in MQ. | Upregulated expression in INFM | Ag/Pb | Kwon and Smith (2022) |
| Vitellogenin (Vg) | Precursor of egg storage protein (vitellin) in MQ | Accumulation in haemolymph of INFM<br>Downregulated expression in INFM<br>Up/Downregulated* expression in INFM | As/Py<br>Ag/Py<br>Aa/Pg | Jahan and Hurd (1998)<br>Ahmed et al (2001)<br>Araujo et al (2011) |
| Vitellogenic Cathepsin B (VCB) | Digests vitellin granules in eggs, rendering amino acids available for the MQ embryo | Up/Downregulated* expression in INF | Aa/Pg | Araujo et al (2011) |

\*Depending on sampling time

**Table 4:** Role of non-immune physiological proteins previously reported as being putatively involved on *Plasmodium* salivary gland survival and onward transmission. Natural mosquito/Plasmodium combinations are in bold. MQ = mosquito, PL = *Plasmodium*, Ag = *Anopheles gambiae*, As = *Anopheles stephensi*, Aa= Aedes aegypti, Pf = *Plasmodium falciparum* (human), Pb = *Plasmodium berguei* (rodent), Pg = *Plasmodium gallinaceum* (avian), INFM = experimentally infected mosquitoes, KDM = knock-down mosquitoes.

| Molecule | Function | Experimental results | MQ/PL combination | References |
| --- | --- | --- | --- | --- |
| Agaphelin | Antihemostatic, facilitates MQ blood feeding. | Upregulated expression in INFM | **Ag/Pf** | Waisberg et al (2014) |
| Anophelin | Antihemostatic, facilitates MQ blood feeding. | Upregulated expression in INFM | As/Pb | Dixit et al (2009) |
| Apyrase (APY) | Anti-hemostatic (blood feeding). Low apyrase has been associated to multiple feeding attempts. When ingested with blood meal prevents blood coagulation in MQ midgut. | Activity lowered in INFM<br>Downregulated expression in INFM<br>Activity lowered in INFM<br>Supplementation increases parasite load | Ae/Pg<br>Ag/Pb<br>Ag/Pb<br>Ag/Pb | Rosignol et al (1984)<br>Rosinski et al (2007)<br>Thiévent et al (2019)<br>Pala et al (2023) |
| Circumsporozoite-protein binding protein (CSPBP) | MQ salivary gland protein. Putative ligand for salivary gland invasion by PL sporozoites | Parasite load decreases in KDM | Ag/Pb | Wang et al (2013) |
| D7 salivary proteins | Salivary gland protein re-ingested by the MQ during the blood meal. Facilitates PL colonization of the midgut (unknown mechanism). Antihemostatic properties. | Parasite load decreases in KDM | Ae/Pg | Martin and Martin (2023) |
| Saglin | Ligand for *Plasmodium* entry into salivary glands<br>Component of MQ saliva, re-ingested by the MQ during the blood meal. Facilitates PL colonization of the midgut. | Downregulation decreases sporozoite load<br>Parasite load decreases in KDM | Ag/Pb, **Ag/Pf**<br>Ac/Pb, **Ac/Pf** | Ghosh et al (2009)<br>Klug et al (2023) |
| SAMSP1 | MQ salivary gland protein. Facilitates sporozoite movement and invasion of host hepatocytes | Parasite load decreases in KNDM | Ag/Pb | Chuang et al (2021) |
| SGS1 | MQ salivary gland protein. Putative ligand for salivary gland invasion by PL sporozoites | Parasite load decreases in KDM | Aa/Pg | Korochina et al (2006) |
| TRIO | MQ salivary gland protein. Facilitates sporozoite movement and invasion of host hepatocytes. | Upregulated expression in INFM<br>Sporozoites from KDM colonize the liver less effectively | Ag/Pf<br>**Ag/Pb** | Marie et al (2014)<br>Chuang et al (2019) |

**Table 5:** Role of non-immune physiological proteins previously reported as being putatively involved on mosquito neuronal function and behaviour. Natural mosquito/Plasmodium combinations are in bold. MQ = mosquito, PL = *Plasmodium*, Ag = *Anopheles gambiae*, Pf = *Plasmodium falciparum* (human), INFM = experimentally infected mosquitoes.

| Molecule | Function | Experimental results | MQ/PL combination | References |
| --- | --- | --- | --- | --- |
| Acetylcholine esterase (Ace) | General neuronal function | Upregulated expression in INFM | **Ag/Pf** | Carr et al (2021) |
| Acetylcholine receptors (AChR) | General neuronal function | Upregulated expression in INFM | **Ag/Pf** | Carr et al (2021) |
| Gamma aminobutyric acid (GABA-A, B) | General neuronal function | Upregulated expression in INFM | **Ag/Pf** | Carr et al (2021) |
| Gustatory receptors (GRs) | Gustatory reception. Modulates sensitivity to sugary food sources | Upregulated expression in INFM<br>Up/Downregulated expression in INFM** | **Ag/Pf**<br>**Ag/Pf** | Carr et al (2021)<br>Hajkazemian et al. (2022) |
| Ionotropic receptors (IRs) | Olfactory responses to host semiochemicals. Host preference and seeking behaviour. | Upregulated expression in INFM<br>Downregulated expression in INFM | **Ag/Pf**<br>**Ag/Pf** | Carr et al (2021)<br>Hajkazemian et al. (2022) |
| Odorant binding and chemosensory proteins (OBPs, CSPs) | Solubility and binding of odorant molecules | Upregulated expression in INFM<br>Up/Downregulated expression in INFM** | **Ag/Pf**<br>**Ag/Pf** | Carr et al (2021)<br>Hajkazemian et al. (2022) |
| Odorant receptors (ORs) | Odour reception | Upregulated expression in INFM<br>Downregulated expression in INFM* | **Ag/Pf**<br>**Ag/Pf** | Carr et al (2021)<br>Hajkazemian et al. (2022) |
| Pickpocket Channel | Gustatory reception | Upregulated expression in INFM | **Ag/Pf** | Carr et al (2021) |
| Transient receptor potential channels (TRPCs and TRPMs) | Gustatory sensitivity | Up/Downregulated expression in INFM | **Ag/Pf** | Carr et al (2021) |
| Transient receptor potential channels (TRP-Waterwitch) | Moisture detection | Downregulated expression in INFM | **Ag/Pf** | Carr et al (2021) |

\* Up-regulated at 7 days post-infection, down-regulated at 14 days post-infection

We then carry out a transcriptomic analysis of malaria-infected mosquitoes, focusing on the physiological and metabolic processes and pathways that have been previously identified as being potentially crucial for the successful development of *Plasmodium* of the mosquito (Tables 1-5). For this purpose, we use the interaction between the most prevalent and widespread avian malaria parasite in Europe, *Plasmodium relictum* (pSGS1), and its main natural vector in the field, the mosquito *Culex pipiens*. *P. relictum* has been at the forefront of research into bird conservation since its accidental introduction to the Hawaiian Islands in the early 20th century, which resulted in the decline and extinction of several native bird species [19,20]. Due to the predicted increase in the occurrence, distribution and intensity of avian malaria associated to the ongoing climate change [21], *P. relictum* is considered an ongoing threat to bird biodiversity and conservation [22]. The IUCN currently classifies *P. relictum* as one of the world’s worst invasive species [23].

Our results provide a comprehensive view of the molecular mechanisms underlying the host’s response to infection and shed light on how *Plasmodium* may exploit mosquito resources to its advantage. We discuss to what extent this may be evidence for parasite adaptive manipulation of its vector or simply the mosquito’s response to the infection. By exploring these interactions, we can gain insights into the co-evolution of *Plasmodium* and their vectors, understanding how selective pressures shape their intricate relationship. This knowledge can also inform the development of new strategies to disrupt these evolutionary adaptations, thereby contributing to more effective control measures against malaria.

## Materials and Methods

### Experimental design

This paper complements a recent paper focused on the immune transcriptome of *Cx. pipiens* mosquitoes infected with *P. relictum* [4]. The aim of the present experiment was to understand how avian malaria infections influence expression patterns of non-immune genes of known relevance to *Plasmodium* development in mosquitoes at four different key stages of *Plasmodium relictum* development within the mosquito [24]: 30Lmin after the blood meal ingestion (30 min) (gametocyte activation and formation of gametes), 8 days post-infection (8 dpi) (peak of oocyst production), 12 dpi (peak sporozoite production) and 22 dpi (end stages of the infection). The full protocol and immune-gene expression data was published in a recent study [4].

Briefly, six canaries, 3 uninfected, 3 infected with *P. relictum* (cytochrome-b lineage pSGS1) were used for the experiments. Ten days later, at the peak of the acute infection stage in blood, the three infected (parasitaemias calculated as % of infected red blood cells: 3.55%, 3.05% and 2.85%) and three control birds were placed individually in an experimental cage with 150 7-day-old female mosquitoes, which had been reared using standard laboratory protocols [25]. Cages were visited 30 min later to take out the bird and any mosquitoes that were not fully gorged. At this point, 10 fully gorged resting mosquitoes were haphazardly sampled from each of the cages. Although mosquitos could have fed from the bird at any time between 1 and 30 min, the samples were lumped for the analyses as a sampling point (henceforth, “30 mpi”). Samples were homogenized with 500 μl of TRIzol LS, and frozen at −80°C for subsequent RNA extraction (one pool of 10 mosquitoes per cage). The rest of the mosquitoes were left in their cages with a source of sugar solution (10%) at our standard insectary conditions (25–27°C, 70% RH). Cages were also supplied with a water container to allow egg laying. On day 8 after the blood meal, 10 mosquitoes were randomly taken from each of the three cages (“8 dpi” sample), homogenized with 500 μl of RNAlater and frozen at −80°C. The procedure was repeated on days 12 (“12 dpi” sample) and 22 (“22 dpi” sample). TRIzol was used for the blood-engorged mosquitoes because bird blood, with its nucleated red blood cells, clogs the filters of the RNA extraction spin columns if they are not first treated in a TRIzol step (see below). To verify the success of the infection, at 8 and 12 dpi, a further sample of 10 mosquitoes per cage was taken and immediately dissected to quantify Plasmodium oocysts in the mosquito gut. All the mosquitos analyzed harboured parasites.

### RNA extraction

All analyses were carried out using pools of 10 mosquitoes (one pool per time point per bird). For the 30 mpi samples, the total volume of the buffer was adjusted to 750 μl; the sample, containing mosquitoes and buffer, was subsequently homogenized using a TissueLyser (Qiagen) equipped with a 5-mm stainless steel bead. The TissueLyser was run for two cycles of 3 min at 30LHz. Phase separation was done according to the TRIzol LS manufacturer’s protocol; the resulting aqueous phase was mixed with one volume of 70% ethanol and placed in a RNeasy Mini spin column. RNA from the 8, 12 and 22 dpi samples was extracted by first transferring the mosquitoes to a new tube together with 600 μl of buffer RLT and a 5-mm stainless steel bead and then homogenized using a TissueLyser. The TissueLyser was run for two cycles of 3 min at 30LHz. Afterwards RNA was extracted using RNeasy Mini spin columns following the manufacturer’s protocol.

The concentration of all RNA samples was measured on a Nanodrop 2000/2000c (Thermo Fisher Scientific). mRNA from each time point was sequenced using an Illumina HiSeq platform at an average of 85 million reads per library (Novogene). We obtained paired-end reads 150 bp in length.

### Data processing

We generated a list of 270 non-immune genes, or gene orthologs, that have been identified, with different degrees of confidence, in the literature as being crucial for *Plasmodium* development within the mosquito (see Table S1). These genes are involved in crucial physiological pathways within the mosquito, and include structural and digestive proteins, hormones, and enzyme involved in the mosquito’s energetic metabolism. At each time point, the expression levels of these genes were compared between infected and control mosquitoes.

Sequence data quality testing was performed using FastQC (version 0.11.8) [26]. Low-quality reads were filtered or trimmed with Trimmomatic (version 0.27) [27]. The resulting files were aligned with Star (version 2.7.9a) [28] by using the *C. quinquefasciatus* genome as reference [29]. Finally, read count per gene was performed using featureCounts (version 2.0.1.1) [30].

### Statistical analyses

All the statistical analyses were carried out with the free statistical software R (R Core Team, 2020) and the free integrated development environment Rstudio (Rstudio Team, 2020). The package DESeq2 (version 1.16.1) [31] was used to estimate the variance– mean dependence in count data from high-throughput sequencing assays and to test for differential expression based on a model using the negative binomial distribution. When testing for significant differences in expression, and to avoid problems arising from sequencing depth, gene length or RNA composition, the count data were first normalized in DESeq2 [32].

To compare gene expression levels between infected and control mosquitoes, we used a differential expression analysis based on the Negative Binomial (a.k.a. Gamma-Poisson) distribution though the function DESeq in the DESeq2 package [32].

## Results and Discussion

In this study, we present a comprehensive transcriptomic analysis of *Plasmodium*-infected mosquitoes, with our findings systematically categorized into five distinct sections based on the potential roles these genes may play in *Plasmodium* fitness within the mosquito vector. The first section delves into genes associated with midgut invasion, providing insights into how *Plasmodium* overcomes the initial barriers to establish infection within the mosquito. The second section focuses on energetic metabolism, highlighting how the parasite utilizes mosquito energetic pathways to support its own energetically demanding development and proliferation. The third section examines genes involved in mosquito life history traits, uncovering how infection influences, in particular, mosquito fecundity. The rationale behind this is that, for horizontally transmitted parasites such as *Plasmodium*, mosquito fecundity is irrelevant to their fitness. Therefore, diverting resources away from egg production towards their own survival and transmission can provide them with an evolutionary advantage, as has been shown in many castrating parasite species [33]. The fourth section explores genes related to salivary gland invasion and onward transmission, elucidating the mechanisms by which *Plasmodium* prepares for transmission to a new host. Finally, the fifth section addresses mosquito biting behavior, offering a deeper understanding of how genetic changes may enhance the likelihood of successful transmission to humans.

### Midgut invasion

Although population reduction occurs at several steps of *Plasmodium’s* life cycle, one of the most drastic bottlenecks takes place between the ingestion of the gametocytes by the mosquito and the production of oocysts on midgut. A study by Gouagna et al, for instance, estimated that in *Plasmodium falciparum*, on average only 0.5% of the gametocytes ingested made it to the oocyst stage [34]. During the first few hours of the infection, the gametocytes need to transform into gametes within the midgut. The search for a mysterious “gametocyte activating factor”, responsible for the exflagellation of male gametocytes to produce eight flagellated male gametes was completed when xanthurenic acid (XA), a byproduct of the mosquito’s kynurenine pathway which metabolizes tryptophan to generate eye-color pigments (ommochromes), was first identified [35–37].

In our transcriptome data, we were able to quantify all 6 enzymes of the kynurenine pathway (Figure 1 and Supplementary Figure S1). The three upstream enzymes that convert tryptophan into the ommochrome precursor 3-hydroxykynurenyne (3HK): tryptophan dioxygenase (TDO), kynurenine formamidase (KFase) and kynurenine mono-oxygenase (KMO) showed a striking pattern of expression, being significantly under-expressed on the first few minutes after the infection, and significantly over-expressed on days 8-12 post infection, during the peak oocyst and sporozoite formation. The fourth enzyme, hydroxykynurenine transaminase (HKT) responsible for transforming 3HK into XA was however, marginally, albeit significantly, under expressed in infected mosquitoes on days 8-12. Of the two enzymes that shunt substrates away from the pathway, PHS did not exhibit any significant differences between infected and uninfected mosquitoes, whereas KAT was marginally (albeit statistically significantly) under expressed day 12 (Figure 1).

**Figure 1:**
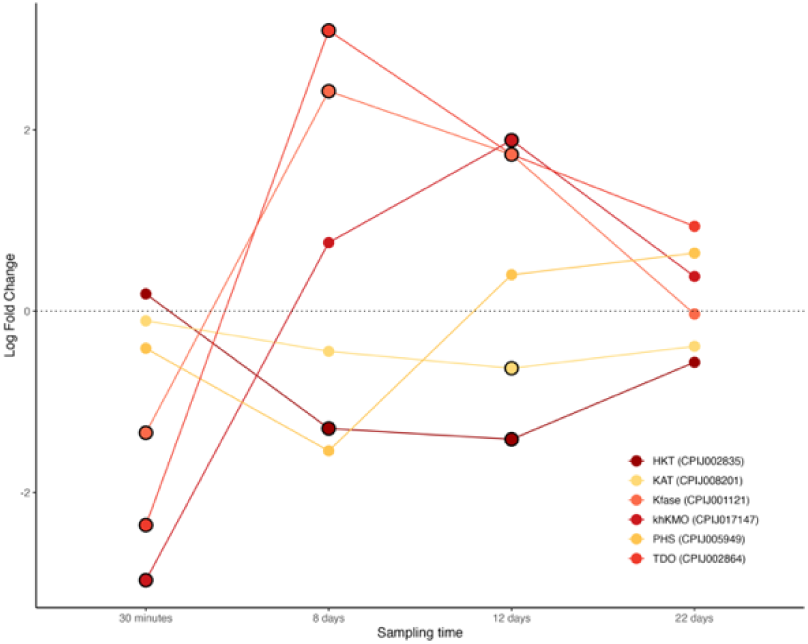
Differential expression pattern of enzymes involved in the kynurenine pathway: Tryptophan-2,3-dioxygenase (TDO), Kynurenine formamidase (KFase), Kynurenine 3-mono-oxygenase (khKMO), 3-hydroxykynurenine transaminase (HKT), Kynurenine aminotransferase (KAT), and Phenoxazinone synthetase (PHS). Differential expression is represented as log-fold change values. Values above and below the dotted line indicate, respectively, increased and decreased expression in infected mosquitoes with respect to non-infected ones. The black circle around each dot indicates that the difference is statistically significant (adjusted p<0.05).

The decreased expression of HKT in infected mosquitoes contrasts with previous results obtained in *Anopheles stephensi*, a human malaria vector, infected with the rodent malaria parasite *Plasmodium berghei* [38] (Table 1). Recent work carried out in this system has uncovered a new role for the kynurenine pathway that does not involve the production of XA as a putative gametocyte activating factor [10]. Knocking down HKT, which transforms 3-hydroxykynurenyne (3HK) into XA (Supplementary Figure S1), significantly increased the number of parasite oocysts in the midgut compared to the controls. The resulting accumulation of 3HK, which has a pro-oxidant activity, impaired the structure of the peritrophic matrix (PM), facilitating its traversal by the ookinetes and the formation of oocysts [10]. Further investigation is necessary to determine whether the overexpression of TDO, KFase, and KMO, along with the under expression of HKT observed in our experiment results in an accumulation of 3HK and a resulting increase in oocyst load in infected *Cx. pipiens* mosquitoes.

The fusion of the gametes leads to a mobile ookinete that needs to traverse the different layers of the midgut to form an oocyst. There is abundant, albeit controversial, evidence that proteolytic enzymes such as trypsins, chymotrypsins, and carboxypeptidases, secreted by the mosquito to digest the blood meal, play a crucial role in the successful formation of oocysts. Baton and Randford-Cartwright suggested that digestive enzymes are largely responsible for the large destruction of ookinetes they observed within the midgut lumen [39]. Other studies, however, established that certain digestive enzymes are, on the contrary, essential for the successful development of *Plasmodium* within the midgut [40–42].

In our dataset, several digestive enzymes were significantly over-expressed in infected mosquitoes (Figure 2A-C). These included aminopeptidase-N, a digestive enzyme that also serves as a ligand for *Plasmodium* in the midgut [43] (Figure 2A), carboxypeptidase-B, which favours oocyst development in the midgut possibly by providing lysine and arginine for the parasite [41,42], and several trypsin and trypsin precursors (Figure 2C). Trypsin is essential for the activation of the *Plasmodium* chitinase, the enzyme that allows the parasite to traverse the peritrophic membrane [11,12,44]. Carboxypeptidase-A, which has been previously shown to favour oocyst development in the midgut through unknown mechanisms [42], was not found in our transcript database. Our results therefore fully agree with previous studies showing an increased expression of these digestive enzymes in infected mosquitoes (Table 1) which may be suggestive of the parasite’s ability to manipulate these enzymes to its own advantage.

**Figure 2:**
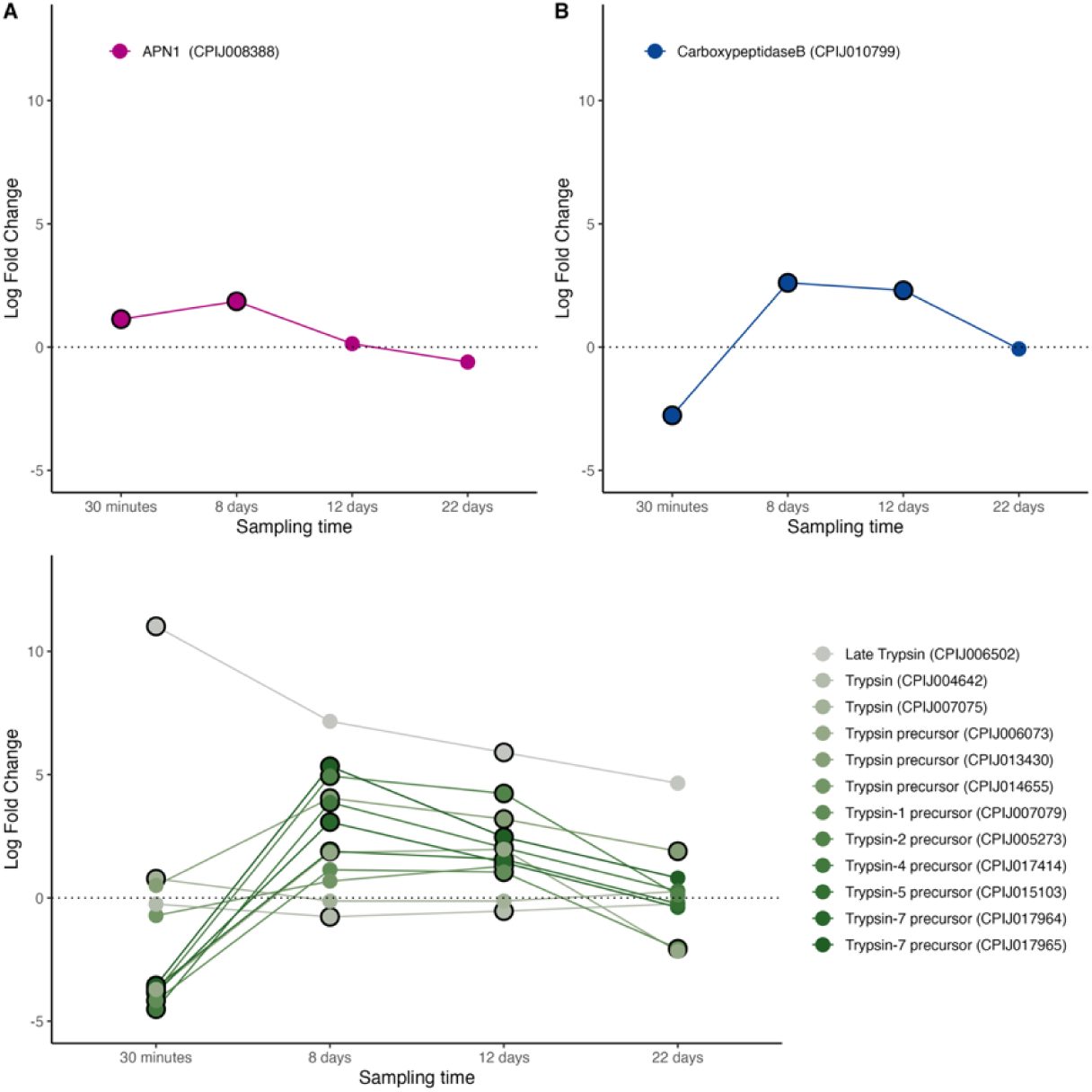
Differential expression pattern of digestive enzymes. A) Aminopeptidase-1, B) Carboxypeptidase-B, and C) Trypsin and Trypsin precursors (see legend), Differential expression is represented as log-fold change values. Values above and below the dotted line indicate, respectively, increased and decreased expression in infected mosquitoes with respect to non-infected ones. The black circle around each dot indicates that the difference is statistically significant (adjusted p<0.05).

Two extracellular structures have been proposed to provide protection to the midgut epithelium in mosquitoes: the peritrophic matrix (PM) and the glycocalyx [45]. These two structures represent a barrier for the invasion of the midgut epithelial cells by *Plasmodium*. The peritrophic matrix is an extracellular sac secreted by the gut epithelial cells, composed of chitin, proteins, and proteoglycans, which completely surrounds the ingested blood meal. Peritrophins are one of the key proteins responsible for the integrity of the PM [46]. None of the peritrophin genes identified from *Anopheles* mosquitoes in VectorBase were found in our transcripts. We did find two genes labelled as PM precursors, one of which (CPIJ010654) was significantly down regulated in infected mosquitoes during the early stages of the infection (30min, Figure 3A). Both precursors where significantly upregulated 8- and 12-days post infection.

**Figure 3.**
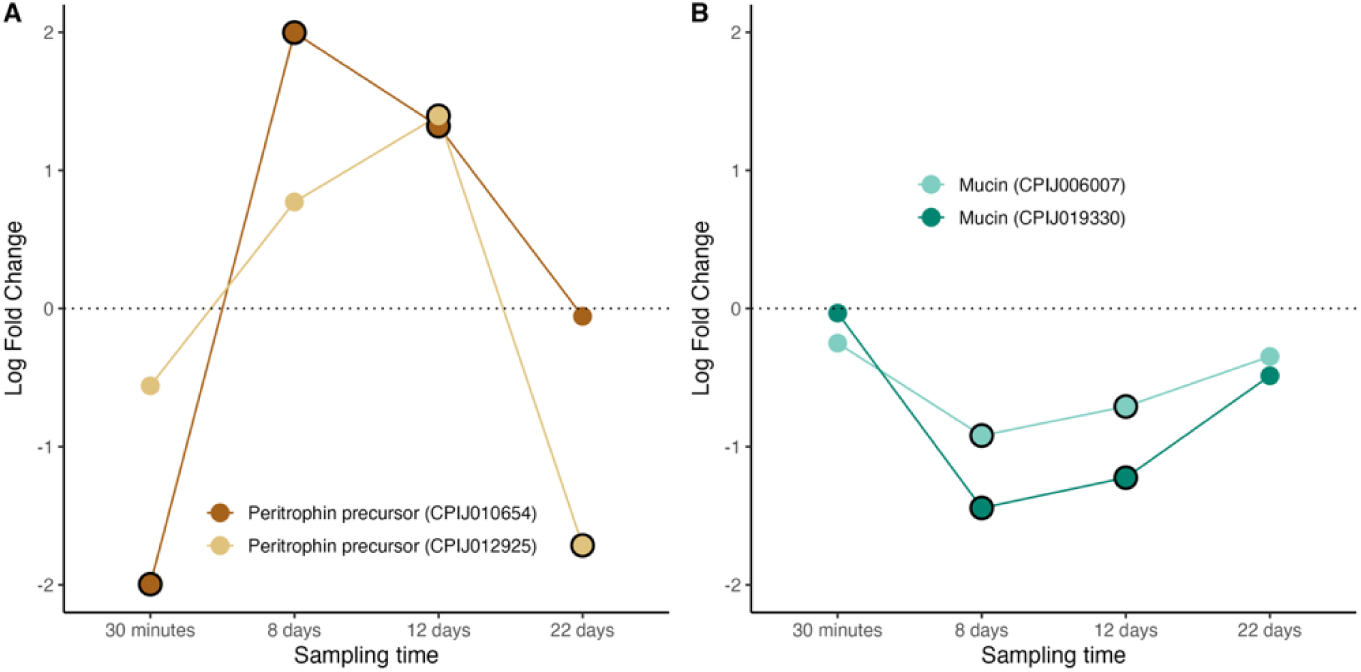
Differential expression pattern of peritrophic matrix (PM) and glycocalyx protein components. A) Peritrophin precursors, and B) Mucins. Differential expression is represented as log-fold change values. Values above and below the dotted line indicate, respectively, increased and decreased expression in infected mosquitoes with respect to non-infected ones. The black circle around each dot indicates that the difference is statistically significant (adjusted p<0.05).

Past the peritrophic matrix, the luminal side of epithelial cells is covered with the carbohydrate-rich glycocalyx. Mucin is one of the key glycoprotein constituents of the glycocalyx. Previous work has established that knocking down the MUC1 gene results in a reduction of oocysts by over 50% [47], and that silencing genes responsible for maintaining the integrity of the mucin barrier, also results in a significant reduction of oocysts in mosquitoes [48,49]. These results suggest that a defective mucin barrier increases the permeability of the midgut to immune effectors, consequently leading to a significant reduction in *Plasmodium* oocysts in the midgut [48]. Interestingly, two mucin genes were identified in our database, both of which were found to highly significantly down-regulated in *Plasmodium*-infected mosquitoes (Figure 3B), suggesting that *Plasmodium* may be able to modulate the expression of these genes in order to maximize its survival in the midgut.

### Energetic metabolism

There is abundant experimental evidence that *Plasmodium* requires extensive carbohydrate and lipid resources in order to fulfil its energetic needs within the mosquito (reviewed in [50]). The most resource-hungry stage of *Plasmodium* within mosquitoes are the oocysts, which constitute veritable DNA-replicating machines, with each oocyst producing thousands of sporozoites. There is increasing evidence that the parasite is heavily dependent on a panel mosquito lipid [13,14] and sugar [15,16] transport proteins as well as enzymatic [17] and hormonal pathways [18] to provide the developing oocysts with these crucial resources.

Several important genes involved in sugar and lipid metabolism were found to be significantly upregulated in infected mosquitoes during the oocyst-formation stage of the *Plasmodium* life cycle. Trehalose transporters are responsible for the transportation of trehalose from the fat body to the haemolymph while glucose transporters mediate the movement of sugars into the cells [16]. Our transcriptome recovered 4 different glucose and 2 different trehalose transporter genes, most of which were significantly over-expressed at 8-12 days post infection, coinciding with the window of oocyst maturation (Figure 4A-B). These results, which agree with previously published studies using other mosquito-*Plasmodium* combinations (see Table 2), could be interpreted in two non-exclusive ways, highlighting the inherent difficulty of interpreting transcriptomic studies as evidence (or lack thereof) of parasite manipulation. On the one hand, it could be a *Plasmodium*-driven strategy whereby the parasite manipulates the expression of these genes to fulfil its energetic means which are known to be consequential [16,51–53]. On the other hand, this could be a mosquito-driven strategy at fulfilling the glucose needs associated to mounting an immune response to the parasite [54].

**Figure 4.**
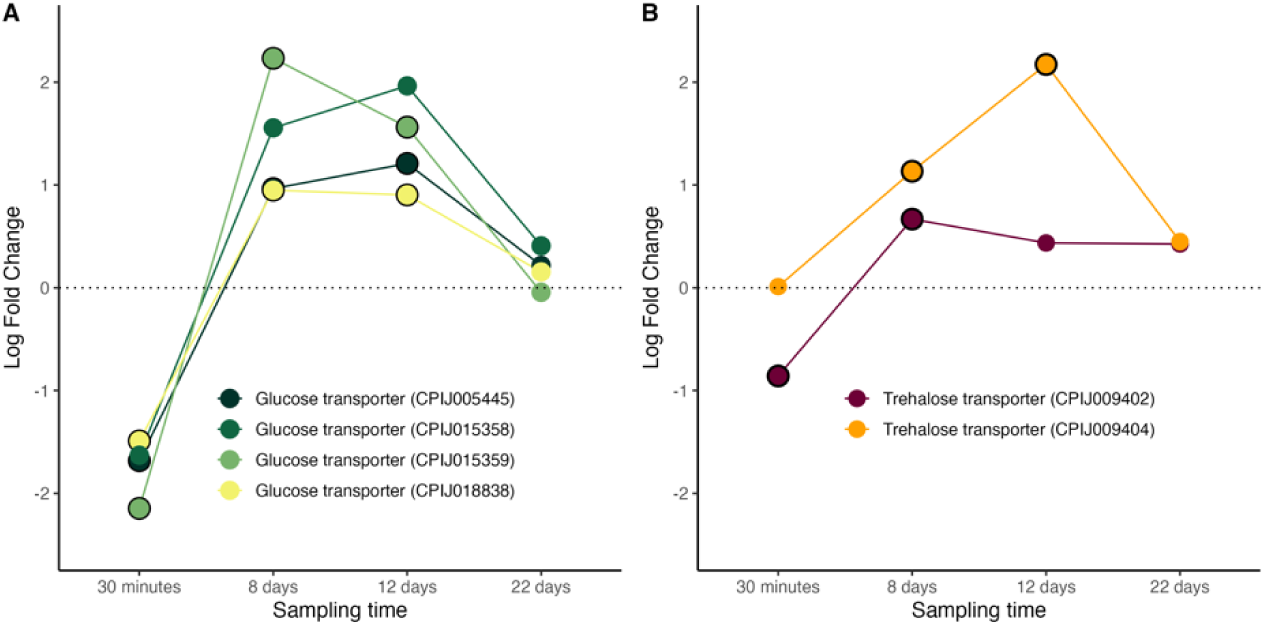
Differential expression pattern of genes involved in sugar transport. (A) glucose transporter genes (B) trehalose transporter genes. Differential expression is represented as log-fold change values. Values above and below the dotted line indicate, respectively, increased and decreased expression in infected mosquitoes with respect to non-infected ones. The black circle around each dot indicates that the difference is statistically significant (adjusted p<0.05).

The main lipid carrier in mosquitoes are lipoproteins called lipophorins [55]. The lipophorin particle consists of two apolipoproteins: apolipoprotein-I, and apolipoprotein-II. There is evidence that apoLp-I and apoLp-II are the product of the same gene and that the two proteins arise from a posttranslational proteolytic processing event [55]. A third apolipophorin, apoLp-III can be found as a lipid-free hemolymph protein that associates with lipophorin during hormone-induced lipid mobilization. Besides its participation in lipid transport, in mosquitoes apoLp-III has also been reported to mediate midgut epithelial defense responses that limit *Plasmodium* infection [56]. We recovered transcripts from one lipophorin receptor (Figure 5A), two apoLp-III (Figure 5B), and one lipophorin precursor (RFABG: retinoid and fatty-acid binding glycoprotein, Figure 5C) all of which were significantly upregulated during the peak oocyst/sporozoite production phase of the infection (days 8-12). These results largely agree with findings in other systems (Table 2).

**Figure 5.**
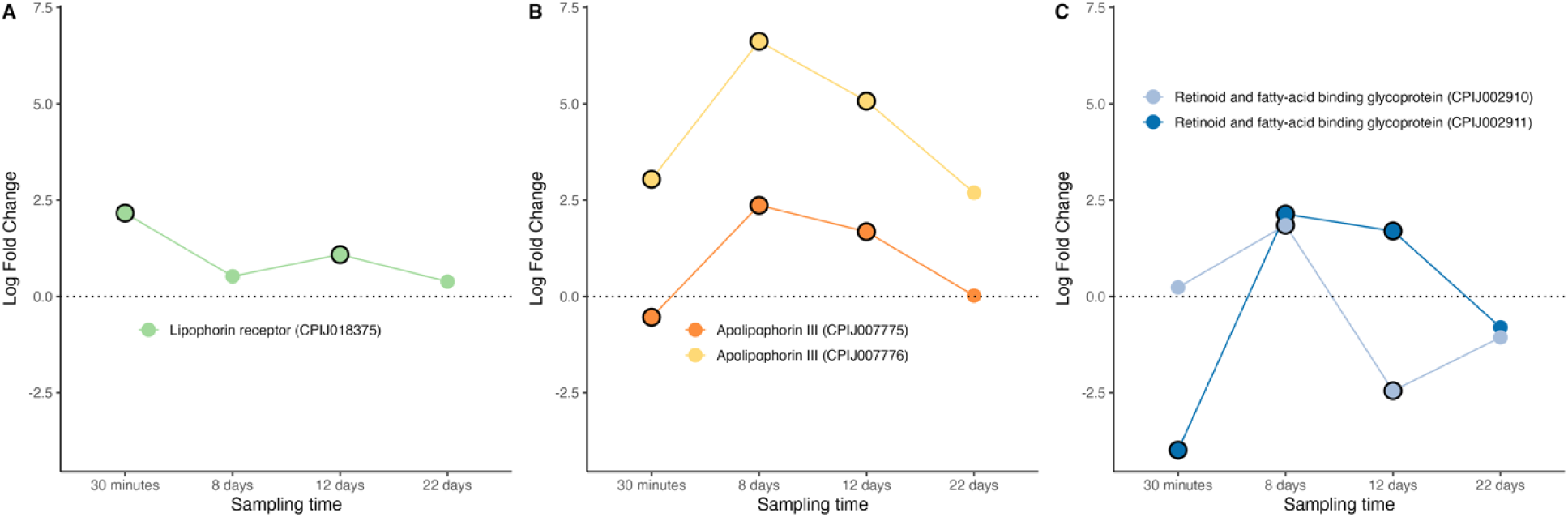
Differential expression pattern of genes involved in lipid transport and metabolism. (A) lipophori receptor, (B) apolipophorin III, (C) retinoid and fatty-acid binding glycoprotein (RFABG). Differential expression is represented as log-fold change values. Values above and below the dotted line indicate, respectively, increased and decreased expression in infected mosquitoes with respect to non-infected ones. The black circle around each dot indicates that the difference is statistically significant (adjusted p<0.05).

AMP-activated protein kinase (AMPK) is a key metabolic energy sensor that regulates total energy stores in many organisms, including mosquitoes [57]. When activated, AMPK inhibits anabolic pathways such as glycogen, protein and fatty acid synthesis and activates catabolic processes that synthesize ATP. AMPK activation also inhibits acetyl CoA carboxylase, crucial in fatty acid synthesis, resulting in decreased levels of cellular lipids [57]. We recovered transcripts from two AMPK genes, one of which showed significant overexpression in infected mosquitoes as compared to uninfected ones on days 8-12 (Figure 6A). This is contrary to previous results showing a diminished AMPK transcripts in the salivary glands of *An. gambiae* mosquitoes infected with *P. falciparum* [58]. Other enzymes implicated in the CoA cycle also showed significant modifications in infected vs uninfected mosquitoes. CoA is synthesized from a precursor called pantothenate via the enzyme pantothenate kinase (PanK). *Plasmodium* parasites cannot synthesize pantothenate, and must therefore uptake it from their mosquito hosts. Previous work has shown that an upregulation of PanK starves the parasite of pantothenate, resulting in significant decreases in parasite numbers [59]. We saw no up or down regulation in either of the two PanK genes we identified in the transcripts of *Cx. pipiens* infected mosquitoes (Figure 6B).

**Figure 6.**
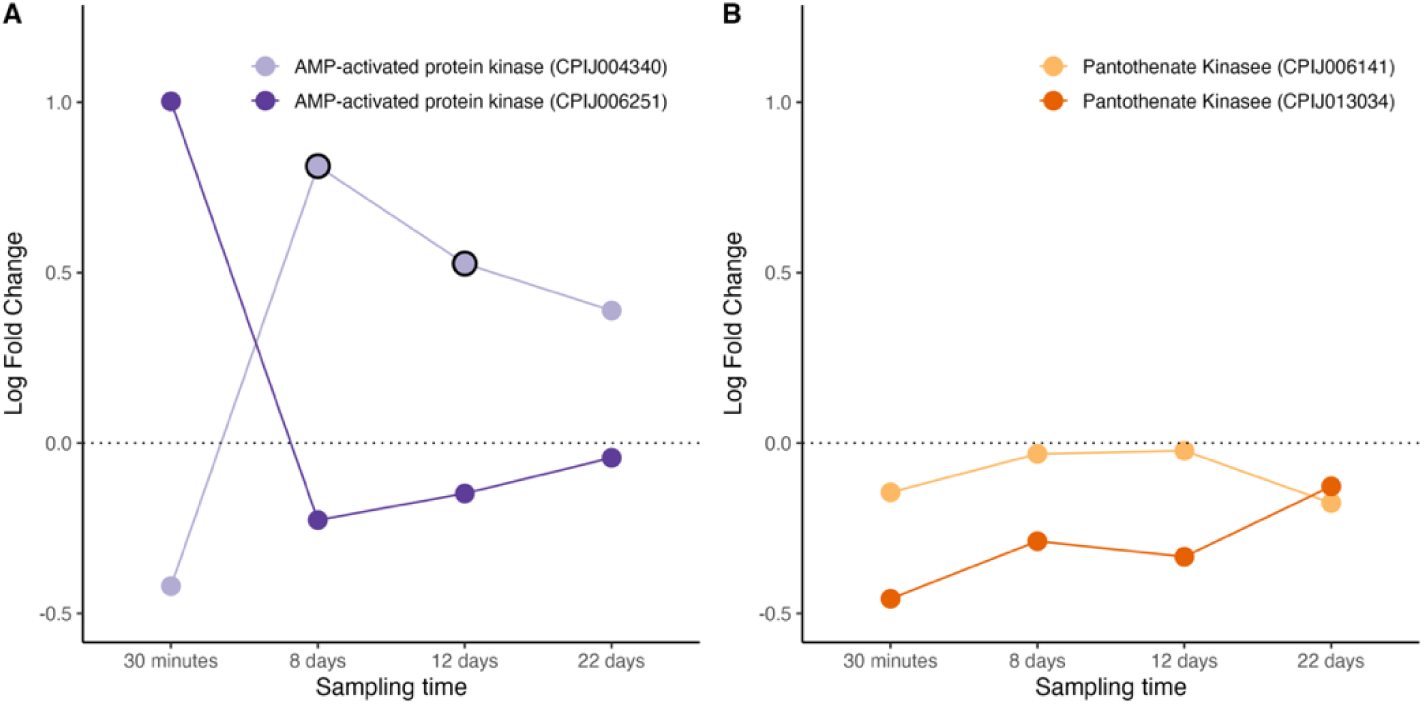
Differential expression pattern of AMP protein kinase (A) and Pantothenate Kinase (B) genes. Differential expression is represented as log-fold change values. Values above and below the dotted line indicate, respectively, increased and decreased expression in infected mosquitoes with respect to non-infected ones. The black circle around each dot indicates that the difference is statistically significant (adjusted p<0.05).

### Fecundity and nutrient recycling

Many parasite species have evolved strategies to reduce the fecundity of their hosts, redirecting resources originally intended for egg production towards their own survival and proliferation [33,60]. Insect eggs are rich in yolk proteins and lipids, which are essential for embryonic development. Since *Plasmodium* transmits only horizontally, mosquito fecundity is of no direct consequence for parasite’s fitness, and studies on mosquito-*Plasmodium* interactions have consistently shown that the parasite frequently reduces mosquito fecundity [61,62]. This has prompted suggestions that, in line with other parasite species, *Plasmodium* may have been selected to either reduce egg production or increase egg resorption in its mosquito host. By redirecting these key resources towards its own metabolic needs and/or towards extending the mosquito lifespan, the parasite would considerably enhance its transmission opportunities. Previous work carried out in our laboratory has shown that a *P. relictum* infection is associated to a significant decrease in fecundity and an increase in mosquito longevity, providing support for this parasite’s ability to reallocate host resources to enhance its own transmission potential [62].

Mosquito ovaries work as conveyor belts where the primary egg follicles are in a pre-vitellogenic state until a blood meal is taken, which triggers a hormonal cascade that initiates the production of yolk protein precursors in the fat body [63]. Once the primary egg follicles are laid, a second burst of reproductive hormones makes the secondary follicles enter a pre-vitellogenic stage where their development is arrested until a second blood meal is taken. At least four key metabolic regulators are involved in egg production in mosquitoes: the TOR (target of rapamycin) nutritional signaling pathway, along with the hormones ecdysone, and juvenile hormone (JH), and insulin-like peptides [64]). TOR is an evolutionary conserved gene that links elevated hemolymph amino acid levels derived from the blood meal to the expression of yolk protein precursors in the fat body. JH stimulates growth and development of the egg follicles and renders them competent to respond to ecdysone [63]. Fat body cells take up ecdysone, convert it to 20-hydroxyecdysone (20E) and use it to activate transcription of vitellogenin genes, the genes encoding the major egg-yolk proteins [65].

In our experiment, mosquitoes laid their first batch of eggs 3-4 days after the blood meal, so mosquitoes sampled on days 8-22 could not have had any fully mature eggs, only eggs in a pre-vitellogenic state (although this was not verified). Despite this, we observed marked differences between infected and uninfected mosquitoes in metabolic pathways involved in egg production. We observed a higher TOR expression in infected mosquitoes immediately after the blood meal, but no significant differences thereafter (Figure 7A). Infected mosquitoes had, however, significantly higher levels of two JH transcripts than their uninfected counterparts particularly on days 8-12 post blood meal (Figure 7B). Although we did not detect any ecdysone transcripts, immediately after the blood meal infected mosquitoes had significantly elevated expression of the ecdysone-response gene E78, a protein that plays a crucial role in egg production in *Drosophila* [66], though has not yet been described in mosquitoes. The expression of this protein falls drastically relative to that of uninfected mosquitoes between days 8-10 post blood meal (Figure 7C). The levels of a precursor of insulin-like peptide (ILPs), which work synergistically with 20E to activate transcription of vitellogenin genes showed an opposite pattern. Expression levels were over 2-fold lower in in infected than in uninfected mosquitoes immediately after the bloodmeal, and were 2-fold higher on days 8-12 (Figure 7D). We also retrieved transcripts from five vitellogenin or putative vitellogenin genes, four of which showed higher expression levels on *Plasmodium*-infected mosquitoes on days 8-12 (Figure 7E). Recently, the enzyme branched-chain aminoacid transferase (BCAT) has been shown to play a crucial role on egg production in mosquitoes [67]. Knocking-down its upstream regulator (miR-276), increases BCAT production which in turn enhances fecundity and decreases *Plasmodium* burden [67]. Interestingly, our results show a drastic down regulation of BCAT in infected, relative to uninfected, mosquitoes on days 8-12 after the blood meal (Figure 7F).

**Figure 7.**
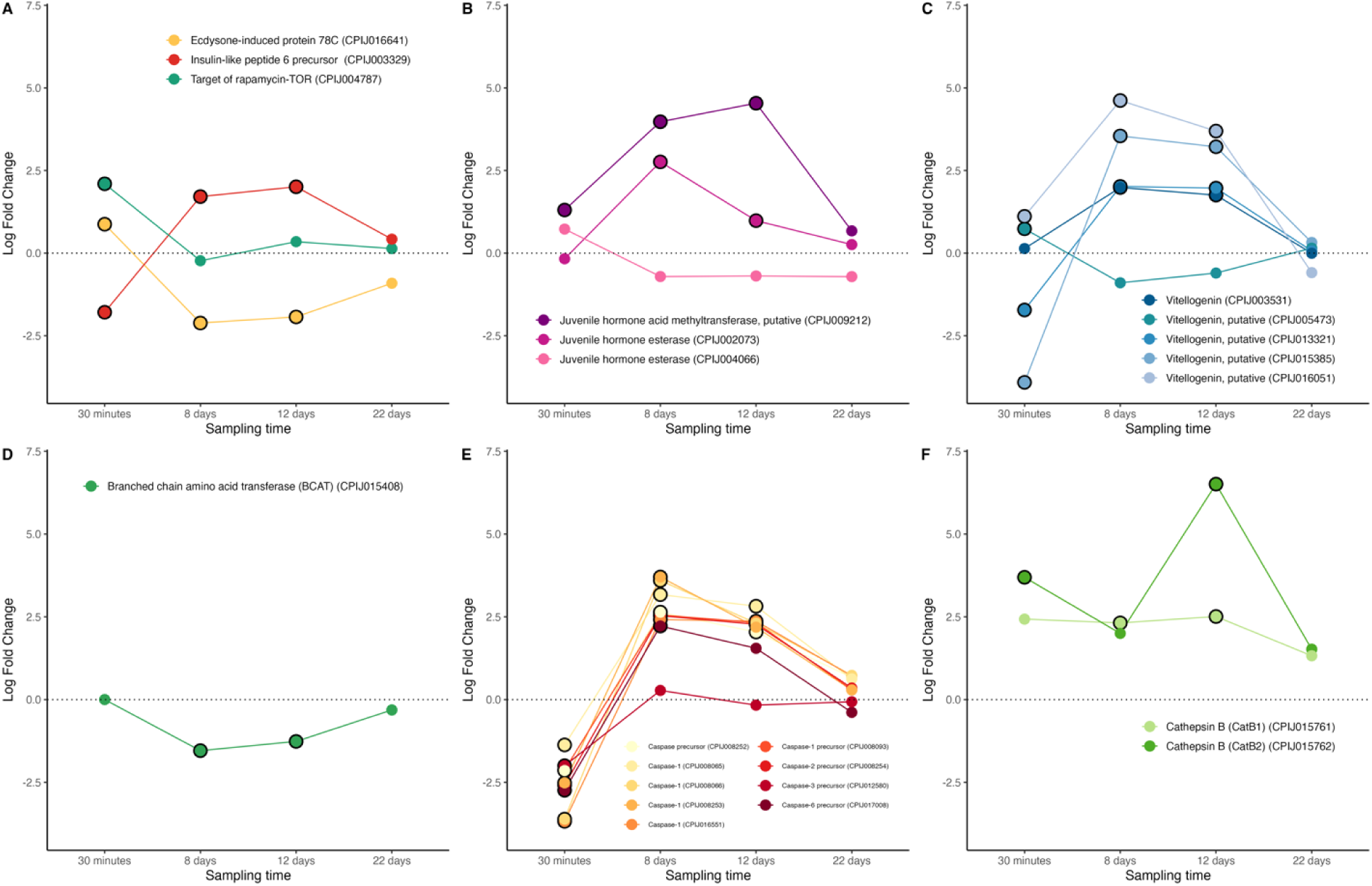
Differential expression pattern of genes involved in egg production and resorption. (A) ecdysone, insulin-like peptide and TOR genes (B) juvenile hormone, (C) vitellogenin, (D) BCAT, (E) Caspases, (F), Cathepsin B. Differential expression is represented as log-fold change values. Values above and below the dotted line indicate, respectively, increased and decreased expression in infected mosquitoes relative to non-infected ones. The blac circle around each dot indicates that the difference is statistically significant (adjusted p<0.05).

Egg resorption is a process that has evolved in many insect species to redirect nutritional resources stored in eggs to other physiological processes of the organism [68]. In mosquitoes placed under stressful conditions, both pre-vitellogenic and vitellogenic follicles may be resorbed [68,69]. Follicle atresia is one of the main factors contributing to malaria-induced fecundity reduction in mosquitoes [61,70]. Two proteolytic enzymes seem to play a central role in egg resorption in mosquitoes: caspases, a family of cysteine proteases that play essential roles in the regulation and execution of apoptosis [61,68] and cathepsin B, a cysteine peptidase that has been shown to be involved in vitellin degradation in mosquitoes [71,72]. We recovered transcripts from 9 different caspases or caspase precursors and two cathepsins, B1 and B2 (Figure 7E and 7F). They all show a pattern of expression that is consistent with a *Plasmodium*-induced trigger of follicular atresia: infected mosquitoes had up to a 4-fold increase in caspase expression between days 8-12 post blood meal and up to a 7-fold increase expression of cathepsin B1 compared to uninfected mosquitoes. These results should, however, be interpreted with caution, considering that caspases, in particular, have been shown to play significant roles in other apoptotic processes associated with *Plasmodium* infection. Most notably, ookinete invasion of the midgut epithelium triggers a caspase-mediated apoptosis of the midgut cells [73] which is why caspases are often considered to be part of the mosquito immune system [74]. Previous work carried out on *P. falciparum* has shown that the parasite is able to evade this arm of the immune system by inhibiting the activity of caspases in the midgut of *An. gambiae* mosquitoes [74,75]. Our findings of a significantly overexpression of caspases at days 8-12 post infection are in stark contrast to these previous results

### Salivary gland invasion and transmission

The salivary glands play a crucial role in *Plasmodium* development because they host the transmissible stages of the parasites (the sporozoites). Sporozoite invasion of the mosquito salivary glands have been shown to require the interaction between *Plasmodium’*s thrombospondinLrelated anonymous protein (TRAP) and a salivary gland receptor in mosquitoes called saglin. Saglin knock-downs resulted in severely decreased sporozoite loads in the salivary glands of mosquitoes [76]. More recently, however, Klug et al. [77] have discovered a different role for saglin in the life cycle of *Plasmodium*. They found that mosquitoes with a knocked-down version of the gene had substantially fewer midgut (oocyst) stages of the parasite than their wild-type counterparts. Their results strongly suggest that saglin is secreted in the saliva and re-ingested during the blood meal, facilitating the colonization of the mosquito midgut through unknown mechanisms [78]. Our results on *P. relictum*-infected *Cx pipiens* mosquitoes show a marginal (albeit statistically significant) saglin under-expression at the beginning of the infection (30min) followed by an over-expression at the peak of oocyst formation in the midgut on day 8 (Figure 8A). However, no significant change in saglin transcripts were detected during the sporozoite stage of the parasite cycle (days 12-22) as would have been expected if the parasite were able to manipulate this salivary gland protein to its advantage.

**Figure 8.**
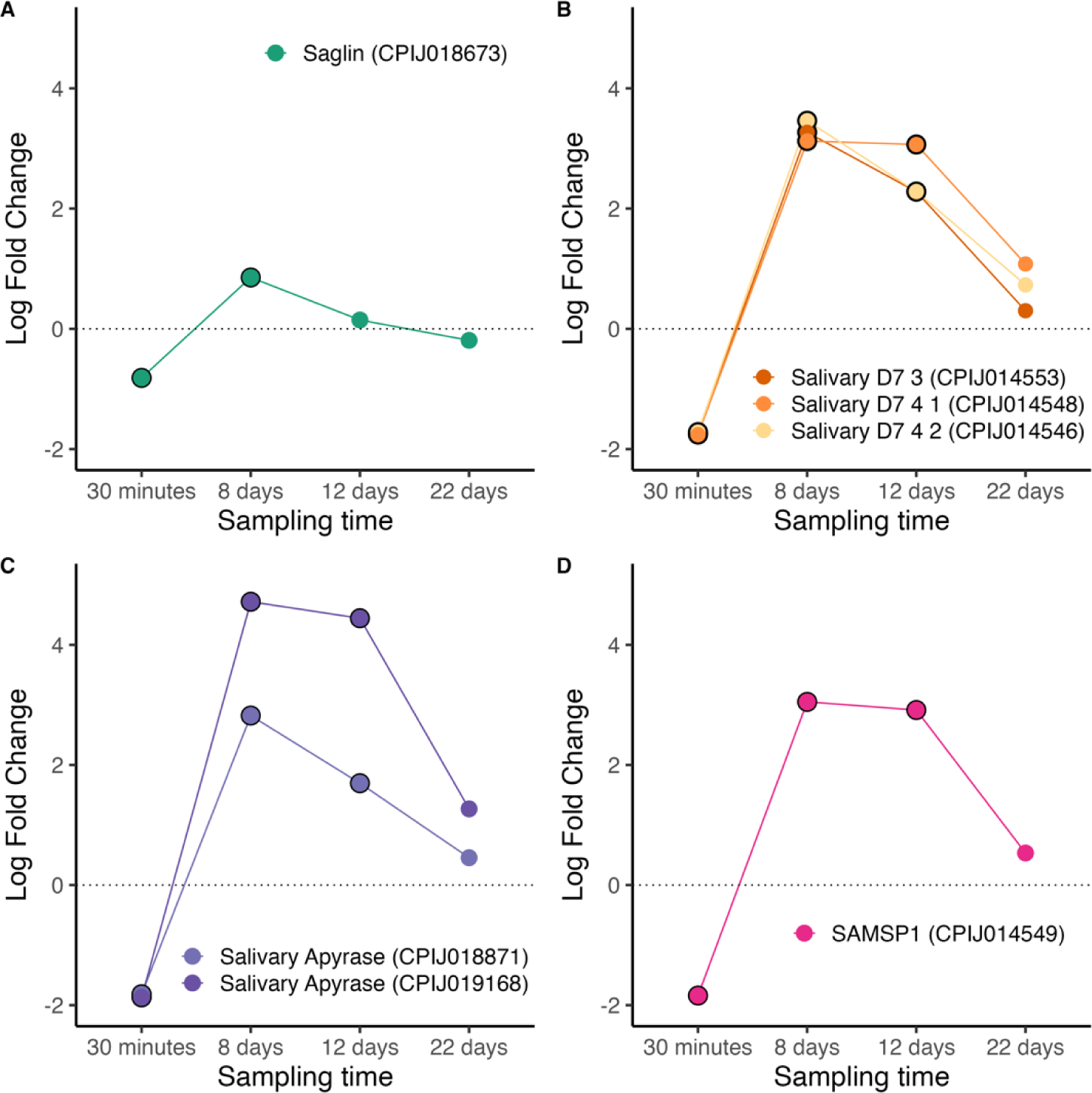
Differential expression pattern of salivary gland proteins. (A) Saglin, (B) Salivary D7 proteins, (C) apyrase (D) SAMSP1. Differential expression is represented as log-fold change values. Values above and below the dotted line indicate, respectively, increased and decreased expression in infected mosquitoes relative to non-infected ones. The black circle around each dot indicates that the difference is statistically significant (adjusted p<0.05).

Previous work has shown that a *Plasmodium* infection has profound effects in several other excreted salivary proteins of mosquitoes [79]. Our transcripts recovered three of these excreted proteins: D7, apyrase and sporozoite-associated mosquito salivary protein (SAMSP1). D7 proteins are among the most abundant components of the mosquito saliva. There are two subfamilies of D7 proteins in mosquitoes: the long-forms (7–30 kDa), and the short-forms (15 to 20 kDa). Although the role of D7 proteins in *Culex* mosquitoes has not yet been entirely elucidated, recent work has shown that two D7 long forms, L1 and L2, have a dual role. On the one hand they have antihemostatic properties that prevent clotting at the site of the mosquito bite, with knock-down mosquitoes having longer probing times than their wild-type counterparts [80]. On the other hand, as with saglin, these salivary gland proteins are re-ingested with the blood meal facilitating the oocyst invasion of the midgut through unknown mechanisms [81]. We obtained transcripts for 3 D7 proteins, all of which showed a significant under-expression in infected mosquitoes at 30 minutes post-blood meal ingestion, followed by a significant overexpression thereafter (Figure 8B).

The first mosquito salivary gland protein to be identified as having a key role in *Plasmodium* transmission was apyrase [82]. Apyrase has been shown to have anti-hemostatic properties, and there are consistent results showing that *Plasmodium* lowers apyrase activity [82,83] and expression levels [84] resulting in prolonged probing times [82]. More recent, though at the time of writing unpublished, results suggest that mosquitoes ingest a substantial amount of apyrase during blood feeding which reduces coagulation in the blood meal by enhancing fibrin degradation and inhibiting platelet aggregation [85]. Supplementation of *Plasmodium* infected blood with apyrase significantly enhanced *Plasmodium* infection in the mosquito midgut [85]. We obtained transcripts from 2 salivary apyrase genes, both of which were significantly under-expressed in infected vs uninfected mosquitoes immediately after the feeding event (30 min) and overexpressed at 8- and 12-days post-infection (Figure 8C). Although the role of SAMPSP1 in mosquitoes has not yet been elucidated, this protein has been recently shown to aid the gliding motility and cell traversal activity of *Plasmodium* sporozoites [86]. In our experiment SAMPSP1 followed the same pattern of expression as D7 and apyrase in infected relative to uninfected mosquitoes: a significant under-expression at 30 minutes followed by significant over-expression during peak sporozoite (day 8) and peak sporozoite (day 12) production (Figure 8D).

### Mosquito behaviour

The purported manipulation of mosquito behavior by *Plasmodium* stands out as one of the most extensively documented instances of parasite manipulation [87]. The mechanistic basis for this manipulation remains, however, largely unexplored. Infected and uninfected mosquitoes have shown differential responses to compounds in host odour using electroantennography [88], and transcriptomic studies carried out in heads or antennae of *An. gambiae* mosquitoes infected with *P. falciparum* sporozoites have shown complex shifts in transcription patterns of key odorant and gustatory genes [58,89].

We detected genes coding for 91 odorant receptors (ORs), 62 odorant-binding (OBPs) and chemosensory proteins (CSPs), and 28 gustatory receptors (GRs, Figure 9). The largest differences between infected and uninfected mosquitoes were observed in OBPs, with 27.5% of genes significantly downregulated shortly after the blood meal (30 minutes) and 27.5-33.7% significantly upregulated during peak oocyst and sporozoite production (days 8-12, Figure 9). A similar, though less pronounced, pattern was found in GRs, where 10.9% of genes were downregulated immediately after the blood meal (30 minutes) and upregulated on days 8 and 12. The smallest changes were seen in ORs, where on days 8-12 the differences between upregulated (2.9-7.4%) and downregulated (0.4-1.5%) genes were less notable. Our results indicate a significant shift towards greater odour and gustatory sensitivity of the mosquito during the transmissible stages of the infection. These results broadly agree with recent transcriptomic studies carried out in heads of *An. gambiae* mosquitoes infected with *P. falciparum* sporozoites [58] and are congruent with studies showing significant shifts in mosquito behaviour during a *Plasmodium* infection. Mosquitoes infected with *Plasmodium* have demonstrated increased attraction to vertebrate hosts, greater persistence in biting, and a higher frequency of feeding when they are carrying the infective sporozoite stage of the parasite [90–92] (but see [93]). These behavioral modifications enhance transmission opportunities for the parasite and have therefore been widely interpreted as being the result of parasite manipulation. [5,92] demonstrated that these shifts in behaviour were not exclusive to *Plasmodium* infections. Mosquitoes infected with *Escherichia coli* exhibited similar shifts in feeding preferences, leading the researchers to conclude that this shift, possibly mediated by insulin-like peptides (ILPs) in the midgut, was not due to parasitic manipulation but rather the mosquito’s immune response to infection [5]. The parasite, they posited, may have evolved to profit from this shift in mosquito physiology.

**Figure 9.**
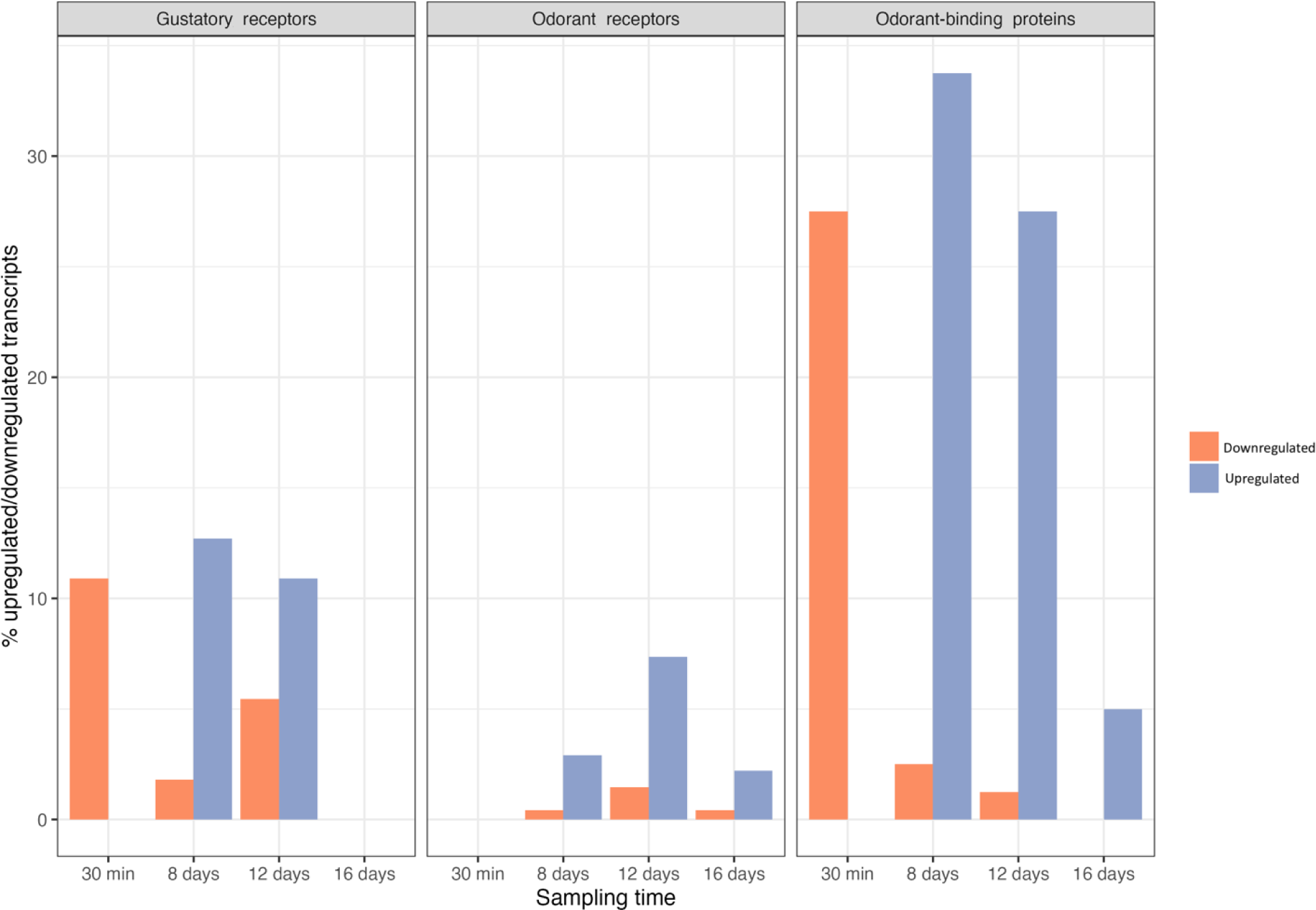
Percentage of gustatory receptors, odorant receptors and odorant-binding protein transcripts that are significantly down (orange) or up (blue) regulated.

## Conclusion

It is evident that a *P. relictum* infection significantly impacts a multitude of physiological pathways in *Cx. pipiens*, in ways that can potentially boost its development and maximize its transmission rates. We observed a significant upregulation of key genes involved in sugar and lipid metabolism, during the crucial oocyst-formation stage, at a time when the parasite critically needs these resources to build thousands of sporozoites, each with its own membrane, haploid genome, cytoplasm and organelles. We also observed significant alterations in key metabolic pathways essential for egg production and resorption, indicating a disrupted reproductive system. These changes provide insights into the mechanisms behind the reduction in fecundity associated with malaria infection in mosquitoes [62,94,95]. This reduction in reproductive output, which in *P. relictum*-infected mosquitoes has been shown to be as high as 40% has been shown to be associated to a significant increase in mosquito longevity a crucial determinant of parasite transmission [62]. Additionally, transcripts for several salivary gland proteins showed significant variations in expression, consistent with previous findings that proteins like apyrase and D7, play crucial roles in enhancing *Plasmodium* transmission. Significant shifts in gene expression related to odorant and gustatory receptors in mosquitoes infected with *Plasmodium*, particularly during the transmissible stages of the infection also align with previous research indicating increased infected mosquito attraction to vertebrate hosts [96].

The difficulty lays in establishing whether these shifts are the result of parasite manipulation or simply the mosquito’s response to the infection, which the parasite may serendipitously exploit to enhance its fitness. From an evolutionary perspective, identifying whether these modifications are driven by the parasite or the vector is a crucial step towards understanding the selective pressures acting on this infectious system. Short of identifying the precise mechanism through which *Plasmodium* may be manipulating its host, something that to our knowledge has not yet been accomplished, one option would be to follow Cator et al’s approach [92,97] and test whether infections with non-vector-borne pathogens yield the same results. An interesting follow-up to this study would thus be to determine which, if any, of the observed transcriptional modifications observed in *Cx pipiens* are absent in mosquitoes infected with *E coli*.

Due to the ongoing climate change, avian malaria is projected to become more frequent. Rising temperatures and changing weather patterns are expanding mosquito habitats, increasing the range of avian malaria and affecting previously unexposed endemic bird species. This shift disrupts ecosystems and threatens biodiversity. Addressing this issue requires a multi-faceted approach, including monitoring vector populations, and developing strategies to block the development of the parasite within the mosquito. The concept of mosquito population replacement, driven by advancements in gene-drive technologies, is emerging as a promising malaria control strategy. This strategy relays on the construction of transgenic mosquitoes that are resistant to *Plasmodium* through a variety of different blocking mechanisms to prevent the parasite from evolving resistance. Although the most widely studied examples of this approach utilize the mosquito’s immune responses, the use of mosquito-derived pathways that facilitate parasite infection and transmission is a promising avenue [98]. Our study uncovers for the first time the specific physiologic pathways that *P. relictum* exploits to thrive within its mosquito vector, providing critical insights for developing targeted interventions to disrupt these pathways and inhibit parasite development.

## Supplementary Figures

**Figure S1:**
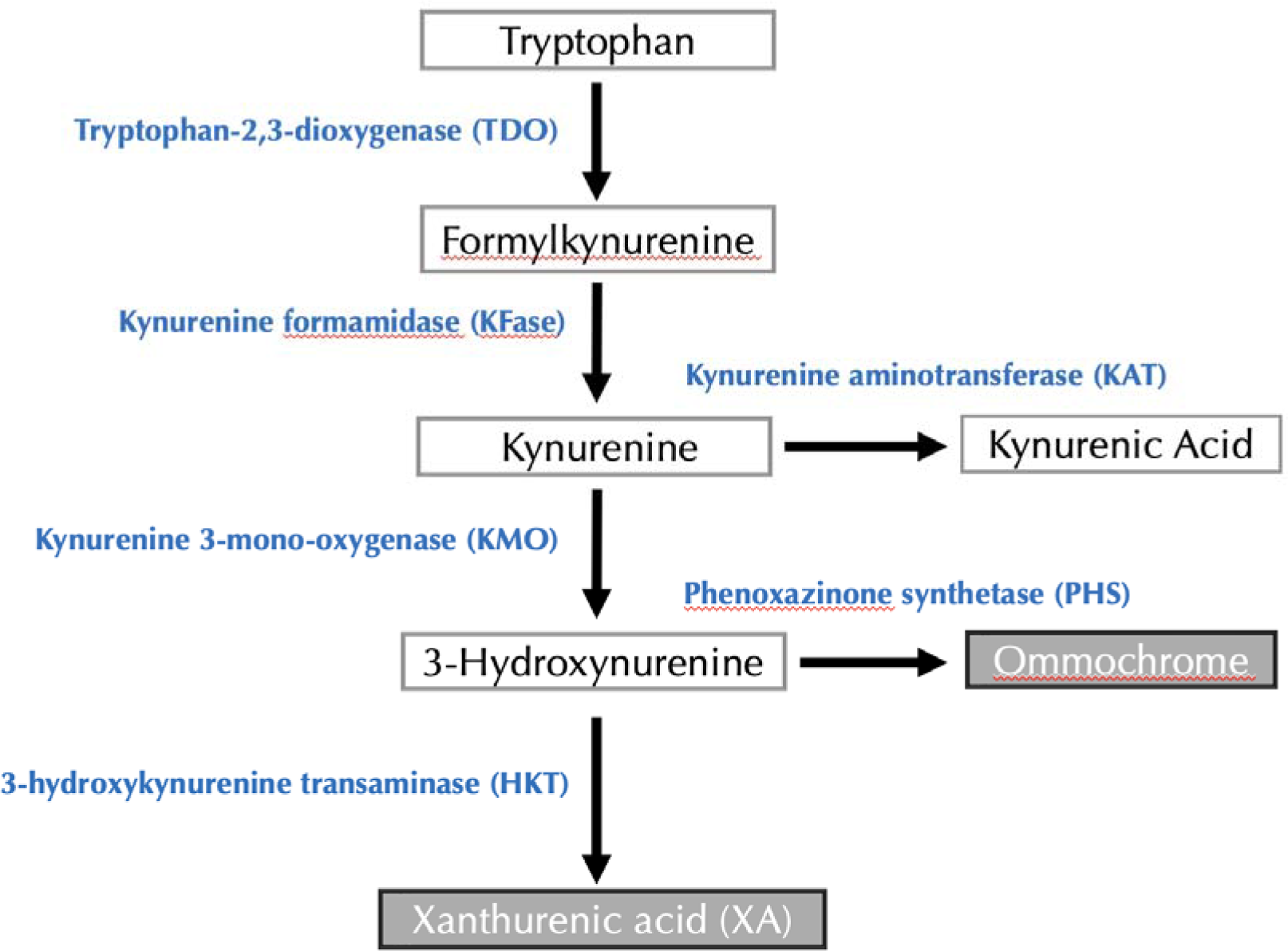
Tryptophan Degradation via the Kynurenine Pathway. The diagram shows the conversion of tryptophan through the kynurenine pathway. Tryptophan is converted to formylkynurenine by TDO, then to kynurenine by KFase. Kynurenine is metabolized to either kynurenic acid via KAT or to 3-hydroxykynurenine by KMO. 3-Hydroxykynurenine is further converted to xanthurenic acid by HKT or to ommochrome by PHS.

